# Kinemomics: spatiotemporal morphodynamic mapping of ventricular kinematic subpopulations in organotypic fetal heart slices

**DOI:** 10.64898/2026.05.19.726402

**Authors:** Gening Dong, Linyang Wu, Junho Choe, Dylan Mostert, Mingkun Wang, Jonathan T. Butcher

## Abstract

Recent advancements in omics technologies have deepened our understanding of complex biological systems and the interplay among gene expression, mechanical cues, and cellular metabolism. However, capturing the spatiotemporal dynamics of live cellular behavior within intact tissue contexts remains a major challenge. Here, we introduce Kinemomics, a framework for spatiotemporal mapping of morphogenetic kinematic states in living tissue. Using a novel fetal cardiac organotypic slice culture system that preserves both beating and morphogenesis, Kinemomics integrates live fluorescent labeling, timelapse imaging, and endpoint immunofluorescence to generate morphodynamic profiles across full-thickness developing myocardium. We validate Kinemomics under controlled Yap-1 signaling perturbations and demonstrate its ability not only to intrinsically decode region- and treatment-specific phenotypes, but also to resolve their spatiotemporal evolution during development and malformation. Kinemomics further reveals localized emergent developmental and malforming fronts that drive system-level behavior, establishing a generalizable platform for investigating computational morphodynamics and cross-scale interactions in complex spatiotemporal systems.

## Main

Recent advances in omics technologies have enabled single-cell phenotypic profiling across multiple molecular levels^1,2^. In parallel, spatial multi-omics approaches have rapidly emerged as powerful means to resolve tissue architecture and cellular heterogeneity^1,3^. However, capturing temporal dynamics underlying biological processes remains challenging. Although single-cell omics studies have modeled cellular differentiation computationally using trajectory inference from reduced-dimensional embeddings^4–7^, these pseudotime analysis relying on predefined gene temporal hierarchies often fail to reconstruct trajectories in physical time within tissue contexts^8,9^.

In developmental biology, cellular kinematics, heterogeneous interactions, and mechanical cues joint to govern embryonic morphogenesis^10–12^, a process of form acquisition not fully captured by genomics and transcriptomics alone^13^. Building on the concept of cellular agency, whereby cells autonomously construct complex tissues rather than rigidly following genetic determinism^14^, modern studies have highlighted dynamic cellular morphing decisions^15–17^. This perspective is further supported by the persistently low proportion of clinical malformations with identified genetic causes^18,19^. Although integration of omics with live imaging has advanced understanding of early morphodynamics^12,20,21^, later-stage processes, especially underlying gestationally survivable but clinically serious congenital defects, remains poorly understood and are largely based on static tissue sections^22,23^. A major challenge is the spatiotemporal quantification of dynamic cellular behaviors in structurally complex and inaccessible tissues, underscoring the need for quantitative frameworks to resolve cellular morphing decisions at clinically relevant stages.

Here, we introduce Kinemomics, a spatiotemporal omics framework built on live timelapse imaging of collective cellular kinematics. Using ventricular myocardium as a model, we resolved layer-specific morphogenetic behaviors underlying trabeculae-compact patterning during development and malformation. We developed an *ex vivo* fetal cardiac organotypic slice culture system that preserves physiological contractility and morphology, enabling longitudinal live imaging and cell tracking within intact four-chamber tissue at previously inaccessible yet clinically relevant developmental stages. Using this platform, we quantitatively mapped mesoscale kinematic behaviors *in situ* under Yap-1 perturbation and demonstrated that local cellular kinematics constitute a morphodynamic fingerprint for high-dimensional subpopulation profiling, enabling delineation of layer-specific phenotypes and tissue-domain boundaries without reliance on RNA sequencing.

By co-mapping live imaging with endpoint immunofluorescence, we linked dynamic cell behaviors to biomolecular activity and identified adaptive phenotypes and emergent kinematic signatures induced by signaling perturbation. Finally, Kinemomics resolved spatiotemporal developmental transitions by identifying time-dependent phenotypic evolution. Together, Kinemomics established a multiscale framework for high-dimensional morphodynamic phenotyping based on live cellular kinematics, enabling enhanced spatiotemporal resolution of cross-scale developmental processes.

## Results

### Organotypic fetal heart slice culture as a physiologically preserved model

Live heart slicing enables *ex vivo* culture while preserving tissue integrity and provides access to four-chamber architecture including full-thickness trabecular and compact myocardium (Fig. 1A, S1A, Supplementary Note 1). Contractile activity recovered within 1–2 hours post-slicing and remained stable for up to 8 hours in culture (Fig. S1B, Video S1). Whole-heart slices exhibited spontaneous conduction without external stimulation (Fig. 2A, Video S2). Cellular calcium dynamics also showed physiologically consistent response under different pharmacological modulations: verapamil reduced beating frequency and prolonged calcium transient duration (CaD50) and off-reaction time (T50 off), epinephrine increased frequency and enhanced calcium transients, and caffeine induced arrhythmic activity (Fig. 2B-C, S1B, Video S3-6), consistent with prior reports^24–29^.

**Figure 1.**
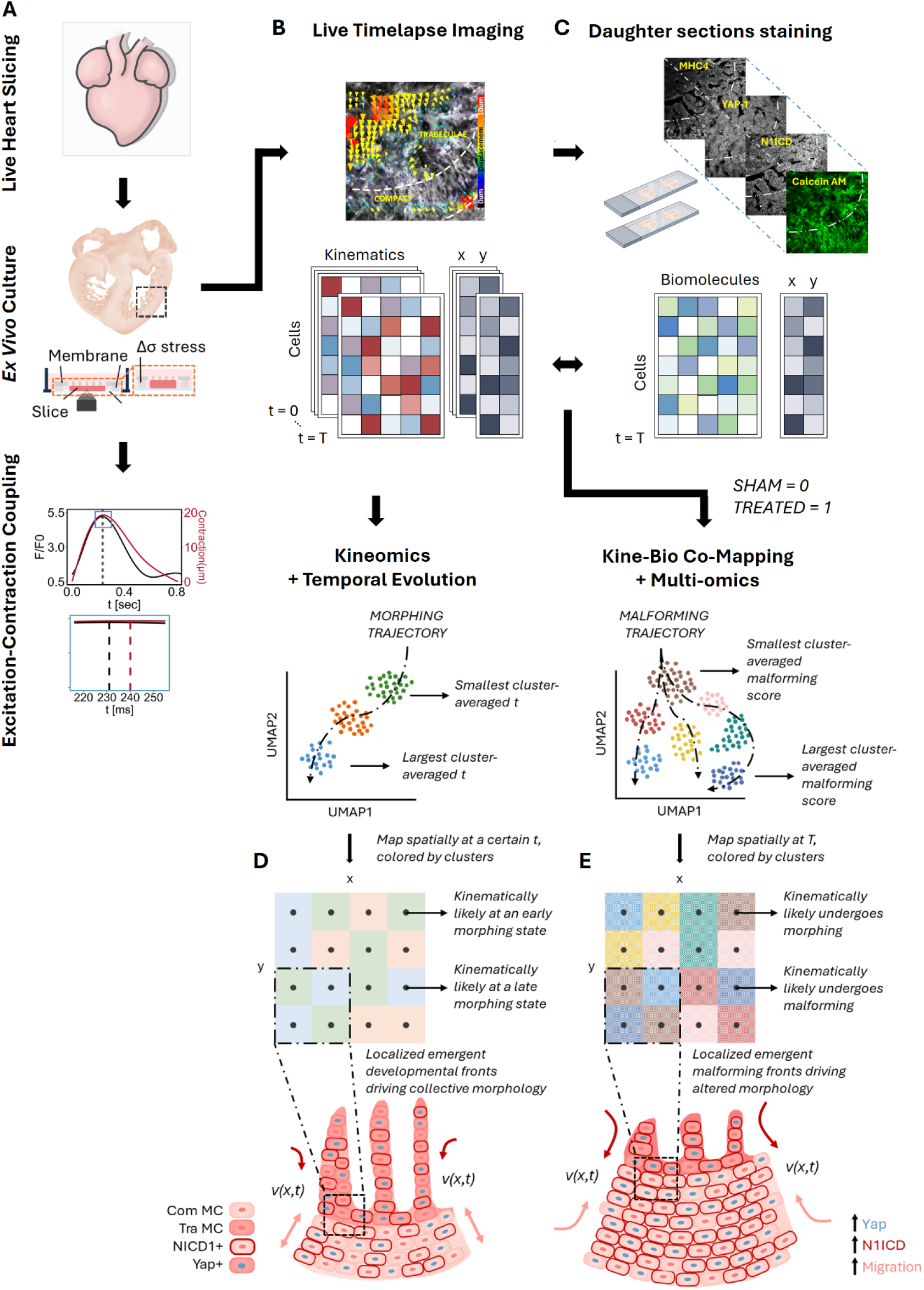
Workflow of spatiotemporal Kinemomics and *ex vivo* slice culture. **A)** Chicken embryonic hearts are isolated, sectioned, and cultured under physiological stress. Cells are labeled with calcium indicator for excitation-contraction coupling analysis and **B)** live cell indicator for timelapse imaging. Local cellular kinematics are resolved spatially and temporally. Kinematic features are extracted and dimensionally reduced for clustering and trajectory inference. **C)** Live heart slices are fixed, and immunofluorescent staining is performed in daughter slices. Biomolecule features are extracted, registered with kinematic feature spatially, and dimensionally reduced for clustering and trajectory inference. Localized high spatial gradient in **D)** morphing progression and **E)** malforming progression was detected as emergent developmental and malforming fronts leading collective tissue morphogenesis and altered remodeling following signaling perturbation, respectively.

**Figure 2.**
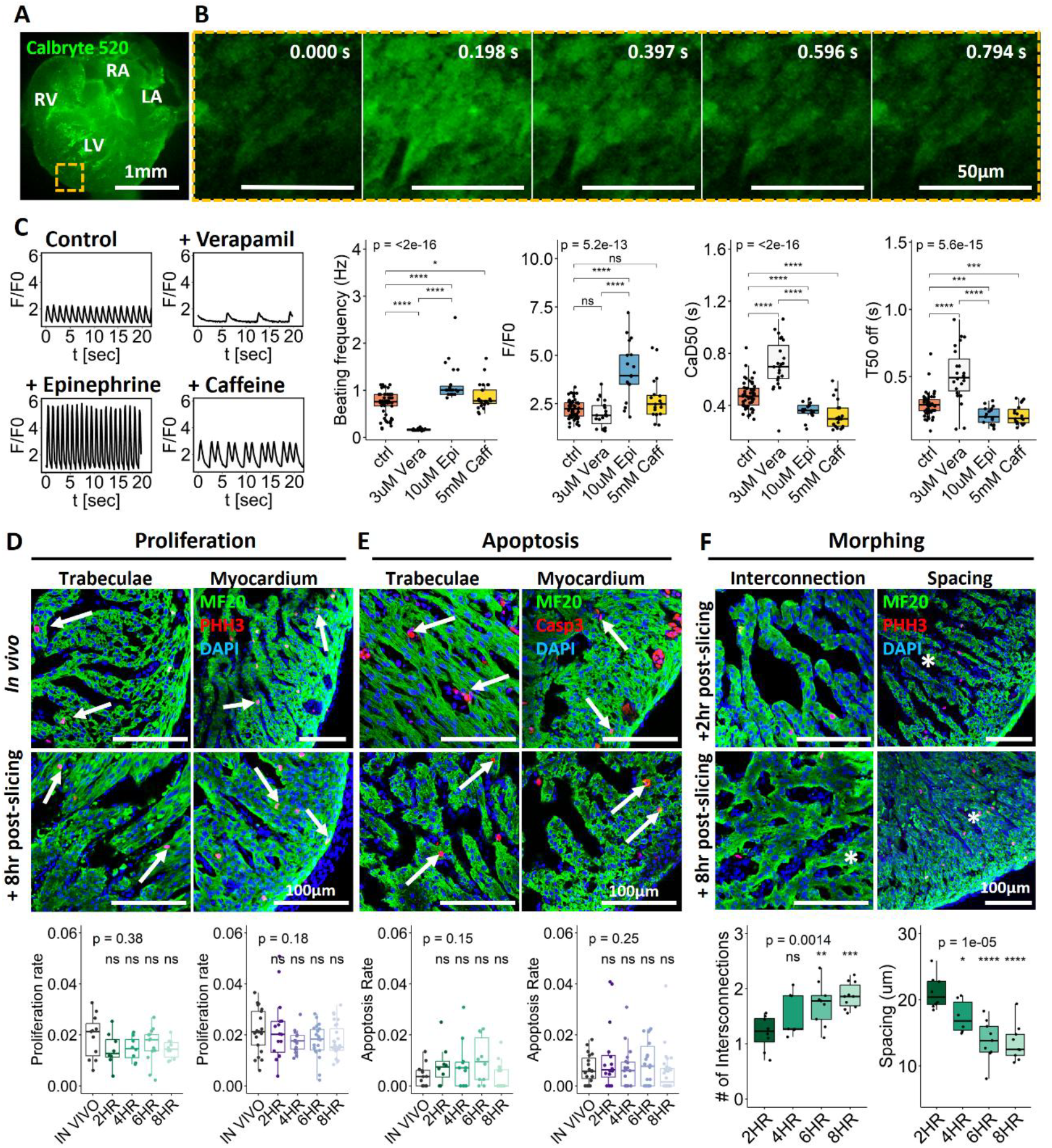
*Ex vivo* heart slice culture as a physiologically relevant dynamic model. 3D live slice culture sustains cardiac contractility, calcium handling and proliferation while morphing under physiological mechanical stress. **A)** Autonomous conduction and **B)** calcium transient in *ex vivo* cultured heart slices. **C)** Individual cell calcium transients (Calbryte 520) tracked during contraction demonstrates physiological calcium handling and pharmacological drug response (e.g. faster and stronger calcium spikes after epinephrine treatment with increased magnitude (F/F0), decreased transient duration (CaD50) and off-reaction time (T50 off); suppression of conduction after verapamil treatment with restricted calcium influx; arrhythmia after caffeine treatment). No difference in **D)** proliferation (pHH3) and **E**) apoptosis (Casp-3) across 8hr culture (white arrows), during which **F)** trabeculae density increased, demonstrating experimental viability and up to 6hr robust morphogenesis without dominant artifacts. *p <= 0.05, **p <= 0.01, ***p <= 0.001,****p <= 0.0001 by ANOVA and t-test with *in vivo* or control as reference group (biological replicates N > 6).

Endpoint assays further showed preserved cardiomyocyte proliferation without increased apoptosis during culture (Fig. 2D-E). Meanwhile, trabecular architecture underwent dynamic remodeling, characterized by increased interconnections and reduced trabecular spacing (Fig. 2F). This was accompanied by compact layer thickening, while trabecular dimensions remained unchanged (Fig. S1C-D), consistent with prior reports at comparable developmental stages^30,31^. Sectioned heart slices also retained mechanosensitive remodeling responses under altered mechanical stress (Fig. S1E). Collectively, these results establish live heart slice culture as a physiologically preserved and artifact-minimized platform for longitudinal morphodynamic analysis.

### Organotypic fetal heart slice as a dynamic testbed to uncover live cellular functional behaviors during ventricular morphogenesis

We next performed timelapse imaging over 3 hours at 10-minute intervals to monitor collective cellular movements within the left ventricular myocardium (Fig. 3A-B, Video S7). Cell tracking and migratory trajectories analyses revealed distinct migratory behaviors between trabeculae and compact myocardium (Fig. 3C, S2A). Although mean migration speed showed no difference between the two layers, trabecular cells exhibited greater net displacement, due to higher migration trace linearity and lower directional change rates relative to compact layer (Fig. 3D, S2B). These findings provide live kinematic evidence supporting earlier static image-based hypothesis that trabeculae growth is driven by directional migration^32^.

**Figure 3.**
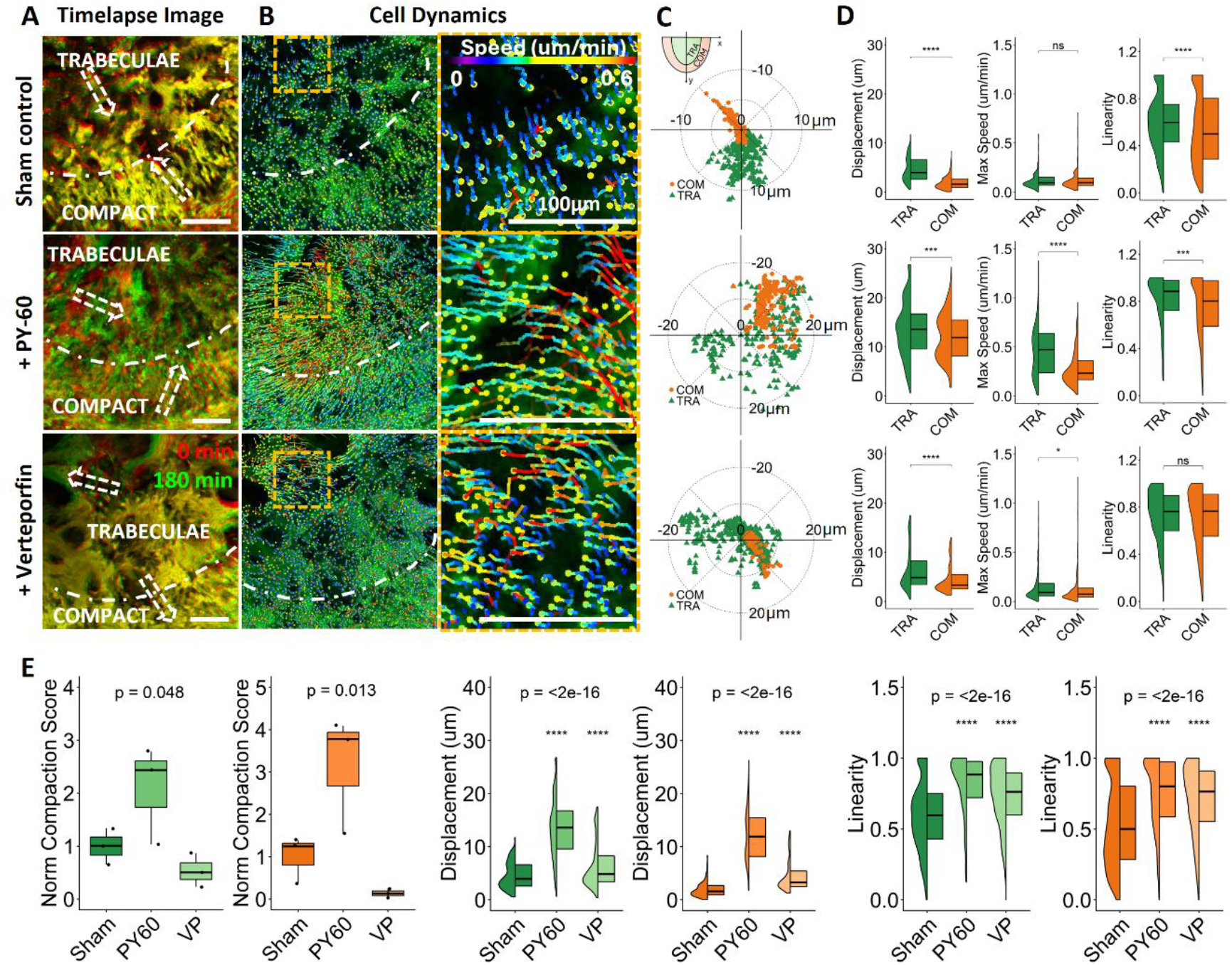
*Ex vivo* heart slice culture as a dynamic testbed to uncover live cellular kinematics during ventricular morphogenesis. **A)** Live slice culture allows for quantitively cell kinematics tracking via live dye labeling (Calcein AM) and timelapse imaging within 6 hr post-slicing in both sham control and YAP-1 perturbed conditions. Live cell tracking recorded distinct myocardial layer remodeling over 3 hr, +2 hr (red) to +5 hr (green) post-slicing, and **B)** local cell migration traces (color traces with the end point as yellow circles). **C)** Local migration directions and displacement were quantitively resolved. Plots showed data from representative samples in sham, + PY-60, and + verteporfin (VP), respectively. **D)** Trabeculae showed larger cellular migration displacement than compact myocardium in all conditions, while cells in compact layer showed a lower migrating trace linearity in sham control and YAP-1 activation. **E)** Cell dynamics was significantly altered by YAP-1 activities. YAP-1 inhibition via VP (10μM) induced local extension in trabecular myocardium with compromised compaction, while YAP-1 activator PY-60 (10μM) caused enhanced compaction with larger displacement. Both YAP-1 activation and inhibition led to higher migrating trace linearity. Normalized compaction score = (proportion of compaction-directed cells)×(mean magnitude of displacement), normalized to that of sham condition. Compaction directions for compact and trabecular layers were defined in supplementary figure S2D. *p <= 0.05, **p <= 0.01, ***p <= 0.001,****p <= 0.0001 determined via ANOVA and t-test with biological replicates N=3 for each condition.

Further, we investigated the role of YAP signaling in regulating cellular morphodynamics. Although the YAP-Hippo pathway is a well-established regulator of cardiomyocyte proliferation, myocardial thickness, and overall heart size^33–38^, the acute cellular dynamics underlying morphological changes remain poorly understood due to limitations of static and endpoint studies. Live slice culture overcomes this constraint by enabling real-time, histology-like visualization of ongoing physiological behaviors within intact tissue. To address this gap, we pharmacologically induced YAP gain- and loss-of-function (GOF/LOF) with PY-60 and verteporfin, respectively. Following YAP activation, compact and trabecular myocardial layers exhibited partially shared morphing direction, resulting in increased myocardial compaction and morphing convergence at the trabecular base (Fig. 3D-E, S2A, Video S8). In contrast, YAP inhibition promoted extension of trabecular tips towards the ventricular lumen, accompanied by outward migration of the compact layer (Fig. 3D-E, S2A, D, Video S9). These timelapse observations provided direct kinematic mechanism underlying prior conclusions inferred from endpoint static imaging, in which cardiomyocyte-specific YAP overactivation produces compact myocardium thickening with reduced trabeculae, whereas YAP LOF results in compact layer thinning and myocardial hypoplasia^33–35^. Importantly, live organotypic slice culture preserves cell-level identities and migratory trajectories, enabling direct resolution of how coordinated cellular behaviors drive tissue remodeling beyond gross organ-level morphological changes. Furthermore, YAP perturbation increased overall tissue displacement, enhanced trajectory linearity, and reduced directional variability across myocardial layers (Fig. 3E, S2C). Directional change rates also trended downward across conditions (Fig. S2E), indicating a shift toward more coordinated and directed morphogenetic states. Together, these findings establish live organotypic slice culture as a platform for resolving spatiotemporal cellular dynamics and kinematic basis underlying tissue morphogenesis and malformation while enabling prediction of subsequent morphogenetic trajectories.

To investigate how functional activity and biomolecular signaling coordinate cell kinematics, we performed calcium imaging followed by immunofluorescent analysis on daughter sections (Fig. 1A, C). Endpoint assays confirmed effective pharmacological modulation, with PY-60 and verteporfin inducing Yap-1 up- and downregulation, respectively (Fig. 4D). Preserved tissue functionality further enabled spatial quantification of excitation–contraction coupling (ECC) (Video S10-11). Yap-1 activation increased calcium transient amplitude, accelerated calcium decay, and shortened contraction time without altering overall ECC efficiency or contractility (Fig. 4A-B, S3A-B). In contrast, Yap-1 inhibition prolonged cellular calcium decay and increased ECC efficiency selectively in the compact layer, while trabecular cells exhibited accelerated off-reaction kinetics (Fig. 4B, S3A-C). This layer-specific functional response was also observed at biomolecular level, where Yap-1 activation increased proliferation in the compact layer but reduced it in trabeculae (Fig. 4C), accompanied by a global upregulation of Notch-1 signaling (Fig. 4E). Consistent with prior studies^35,39^, these findings highlight the capacity of the slice culture system to resolve layer-specific functional and molecular responses within an intact tissue context.

**Figure 4.**
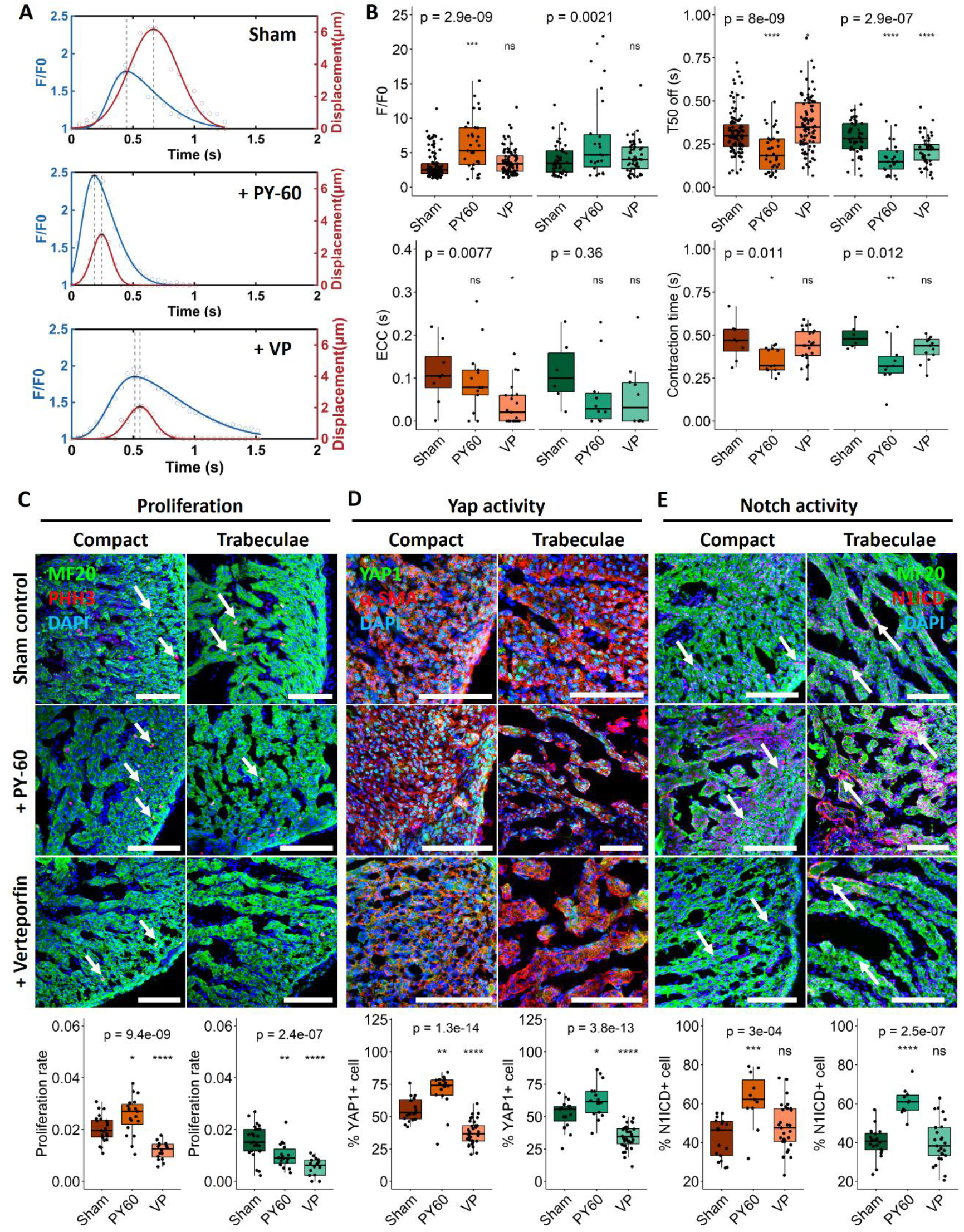
Excitation-contraction coupling in *ex vivo* heart slice and endpoint assay. **A)** Autonomous calcium transient and contractile displacement coupling in *ex vivo* cultured heart slices under different pharmacological agent treatments to alter Yap-1 activity. **B)** YAP activation led to increased magnitude (F/F0), faster decay (T50 off), and decreased contraction time. YAP inhibition showed matured phenotype in the compact layer only with slower decay, higher coupling efficiency (ECC). Endpoint assay in **C)** proliferation, **D)** Yap-1, and **E)** Notch-1 activity. Yap-1 activation led to increased proliferation in compact layer only, along with increased Notch-1 activation. *p <= 0.05, **p <= 0.01, ***p <= 0.001,****p <= 0.0001 by ANOVA and t-test with sham control as reference group (biological replicates N > 6).

### Kinemomics resolves myocardial layer-specific sub-phenotypes

Although embryonic trabecular and compact myocardium exhibit significant architectural and morphological differences, they remain transcriptionally similar with relatively few layer-specific genes reported^40–42^. Consequently, spatial transcriptomics studies have primarily distinguished and subclustered these populations based on gradient enrichment of limited marker genes and their spatial positioning along the ventricular wall^43–45^. Given the substantial differences in morphing directionality and magnitude observed across myocardial layers and signaling perturbations, we next asked whether cellular kinematics could provide a robust morphodynamic fingerprint for refined sub-phenotyping.

To test this, we performed dimensionality reduction and Gaussian mixture model clustering using cell kinematics derived from 3-hour live tracking datasets and validated against manually segmented ground truth (Fig. 1B, 5A-B). Kinemomics identified clusters corresponding to myocardial layer-specific phenotypes (C1, C4 for trabeculae, and C5 for compact in Fig. 5A-C). It also detected hybrid clusters exhibiting intermixing of compact and trabecular cells (Fig. 5A-C), which localized near the interface between manually annotated compact and trabecular layers when mapped back to native spatial coordinates (Fig. 5B). Notably, these interface-associated clusters delineated a clear tissue-domain boundary, suggesting the presence of an interfacial mixing zone between distinct myocardial layers intrinsically decoded by cell kinematics alone. Together, these findings demonstrated that Kinemomics can discriminate sub-populations within distinct myocardial layers and clearly define tissue-domain boundaries, providing informative morphodynamic signatures.

**Figure 5.**
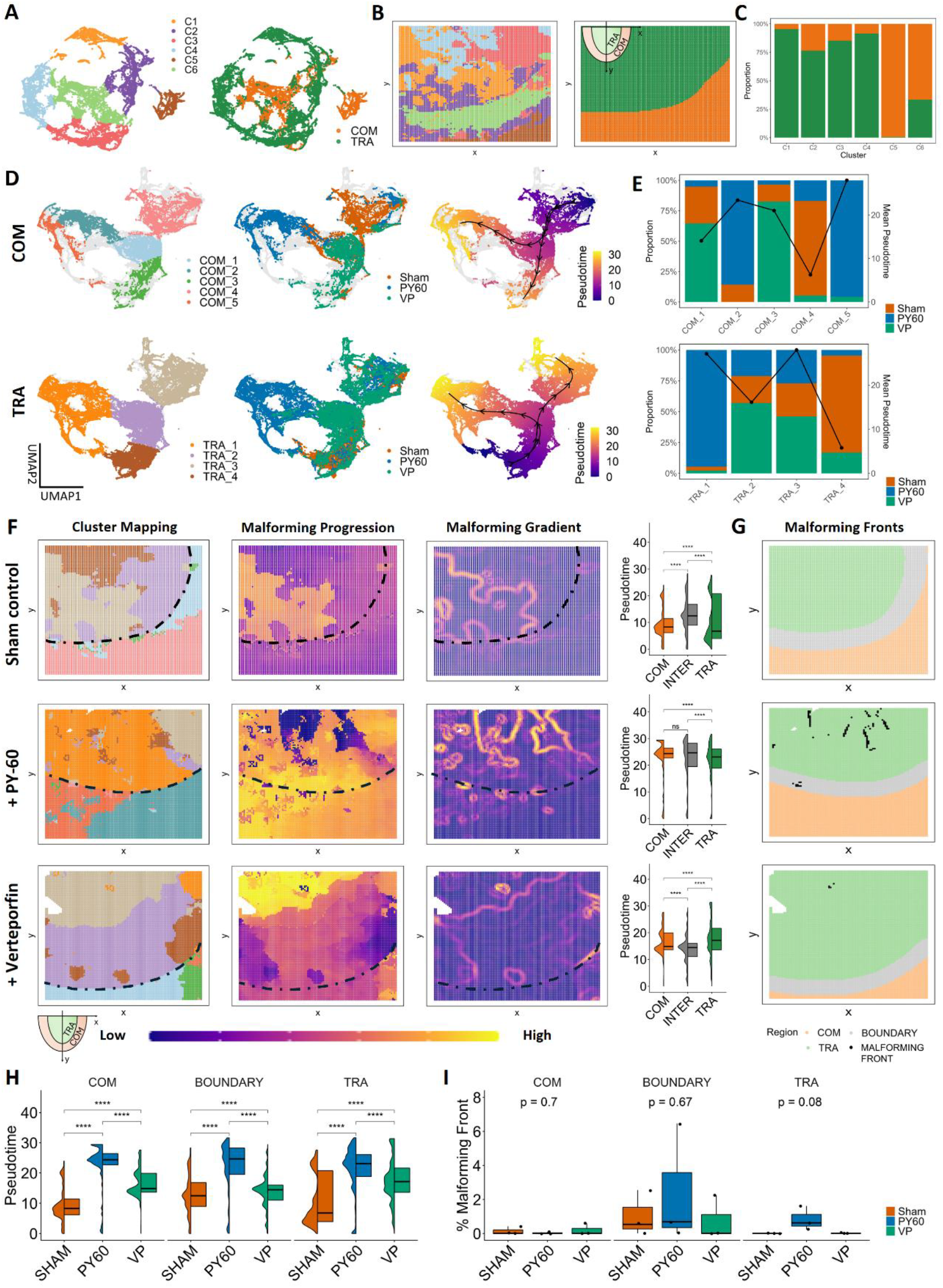
Live timelapse imaging derived local cellular kinematics and endpoint assay enabled kine-bio co-mapping based multi-omics. **A)** High-dimensional omics approaches based on local cellular kinematics is sufficient to identify compact versus trabecular phenotypes. **B)** Consensus spatial mapping of Kinemomics determined clusters versus manually determined regions. **C)** Stacked bar plots showing the cluster fraction of distinct myocardial layers. **D)** UMAP of heart slices in untreated and treated conditions based on kine-bio co-mapping derived features and trajectory inference. **E)** Stacked bar plots showing the cluster fraction of different treatments, overlayered with cluster-wise pseudotime. **F)** Spatial mapping onto original cellular coordinates of multi-omics determined clusters (first column), malforming progression represented by pseudotime (second column), spatial gradient of malforming progression (third column), along with corresponding violin-box plots for region-wise quantification. **G)** Spatial mapping onto original cellular coordinates of malforming fronts, identified by peak malforming gradients (threshold = 0.45). **H-I)** Violin-box plots of region-wise quantification across treatments for pseudotime and malforming front proportion. *p <= 0.05, **p <= 0.01, ***p <= 0.001,****p <= 0.0001 determined via pairwise Mann–Whitney U test or ANOVA with biological replicates N=3 for each condition.

We next extended Kinemomics to datasets subjected to YAP perturbation. Perturbed tissue exhibited reduced kinematic separability between compact and trabecular myocardium (Fig. 3C, S4B, E). Nevertheless, Kinemomics identified compact-enriched (C3) and trabeculae-enriched (C6) phenotypes in YAP GOF samples (Fig. S4A-C), as well as trabeculae-enriched phenotype (C3) in YAP LOF samples (Fig. S4D-F), highlighting the ability of Kinemomics to intrinsically resolve tissue-domain-specific kinematic states. Collectively, these findings establish cellular kinematics as a powerful morphodynamic signature, complementary to transcriptional gradient, for defining tissue-domain boundaries.

### Kinemomics integrated with biomolecular signaling resolves phenotype response to signaling perturbations

After demonstrating Kinemomics can discriminate compact and trabecular myocardium, we extended the framework by co-mapping live imaging-derived kinematic features with immunofluorescence-based biomolecular activity profiles, thereby enriching the morphodynamic feature space (Fig. 1C). We next asked whether this integrated Kine-Bio Multi-Omics framework could resolve malforming progression both kinematically and biologically.

Pooling samples across all YAP conditions, we performed dimensionality reduction and clustering using kinematic features extracted from 3-hour live imaging together with co-mapped endpoint biomolecular assays from matched daughter slices (Fig. S5A-B). Kine-Bio Multi-Omics identified treatment-enriched phenotypic clusters across myocardial layers (Fig. 5D-E, S5C). Sham-enriched clusters exhibited coordinated myocardial compaction (COM_4, TRA_4) with moderate chamber and lumen expansion (COM_1, TRA_2) (Fig. 5E, S6A). YAP activation-enriched clusters displayed distinct morphogenetic behaviors, including compact layer circumferential compaction (COM_2, COM_5) and markedly accelerated trabecular compaction (TRA_1). In contrast, YAP inhibition-dominated clusters showed pronounced compact layer-driven chamber expansion (COM_1, COM_3) and trabeculae extension toward the lumen (TRA_3) (Fig. 5E, Fig. S6A). These findings identify YAP signaling as a key regulator of myocardial compaction, while trabecular extension and chamber expansion likely involve additional pathways that become dominant upon YAP inhibition. Hybrid phenotypes were also observed, suggesting shared morphodynamic features across treatment conditions, potentially reflecting differential YAP sensitivity across tissue domains and relatively short treatment duration. Together, these results demonstrate that Kinemomics integrated with biomolecular profiles resolves treatment-specific phenotypes and uncover emergent malforming signatures under signaling perturbation.

### Kinemomics phenotype detects emergent malforming fronts following signaling perturbations

To further characterize malforming progression, we performed trajectory inference using cluster-wise malforming scores to generate pseudotime trajectories reflecting progressive morphological alteration (Fig. 5D, F, S5D). Trajectories were validated by a strong regression fit and a significantly positive association between pseudotime and the fraction of treated cells per cluster, indicating biological consistency (Fig. S5E). Given the intermixing of compact and trabecular phenotypes near the tissue boundary, we defined an intermediate boundary zone to extend spatial analysis (Fig. S4G). Regional comparisons revealed that YAP activation elevated malforming progression in the compact and intermediate regions, whereas YAP inhibition induced greater morphological alterations within the trabeculae (Fig. 5F). Overall, PY-60 treatment induced more extensive morphogenetic alterations across the full-thickness myocardium than verteporfin (Fig. 5H).

Spatial mapping of clusters and malforming progression onto native cellular coordinates revealed pronounced spatial discontinuities, prompting us to investigate whether Kine-Bio Multi-Omics could identify localized malforming fronts guiding collective tissue remodeling. We therefore quantified spatial gradients of malforming progression to detect localized sharp spatial variations (Fig. 5F, S6B), an approach commonly used to delineate interfaces in physical and geological systems^46–49^, and biochemical gradients governing embryogenesis^50,51^. We hypothesized that steep morphodynamic gradients represent localized drivers of collective cellular behaviors, with peak malforming gradients indicating malforming fronts. Intriguingly, the intermediate boundary zone contained the highest fraction of malforming fronts (Fig. 5G, 5I, S6B), suggesting pronounced spatial heterogeneity in morphological behaviors within the interfacial region between myocardial layers. Furthermore, YAP activation increased the incidence of malforming fronts in the trabeculae while simultaneously inducing pronounced morphological alterations in the compact layer (Fig. 5F, H), suggesting that YAP activation drives global morphogenetic alteration in the compact layer, whereas trabeculae retain localized heterogeneity with partially preserved sham-like morphogenetic signatures. Consistently, Kinemomics clustering revealed a higher proportion of PY-60 phenotypes intermixed with sham and verteporfin-associated phenotypes in the trabeculae (TRA_2, TRA_3; Fig. 5E). Collectively, these findings highlight that Kine-Bio Multi-Omics can resolve treatment-specific phenotypes while identifying localized emergent malforming fronts.

### Kinemomics resolved time-dependent phenotype evolution during myocardial remodeling

Extending beyond spatial kinematic analysis, we next examined time-resolved morphodynamics over a 3-hour live imaging period. We aimed to demonstrate that Kinemomics can resolve not only distinct phenotypes spatially but also spatiotemporal phenotypic evolution. Cellular kinematic features were extracted at 30-minute intervals and reconstructed into sample-specific morphogenetic profiles (see Methods), followed by dimensionality reduction and clustering for each sample. Spatiotemporal Kinemomics identified sub-phenotypes that emerged at distinct time points of live tracking (Fig. S7-10). When mapped back to their original spatial coordinates, these sub-phenotypes revealed localized temporal transitions (Fig. S7-10), indicating dynamic shifts in morphogenetic behavior over development.

To quantify these morphodynamic transitions, we performed trajectory inference and designated clusters with the shortest mean tracking duration as start clusters. The resulting pseudotime trajectories therefore provided a continuous metric of morphing progression (Fig. 1B, 6A). Further, we constructed morphing-state matrices including morphing speed, temporal heterogeneity, and morphing activity derived from UMAP-based trajectories (see Methods), enabling systematic characterization of spatiotemporal dynamics (Fig. 6B, S11). Under sham condition, trabeculae displayed the highest morphing speed, the intermediate boundary zone showed greater temporal heterogeneity, and the compact myocardium exhibited the lowest morphing activity, indicating that trabeculae cells undergo more dynamic remodeling (Fig. 6C).

**Figure 6.**
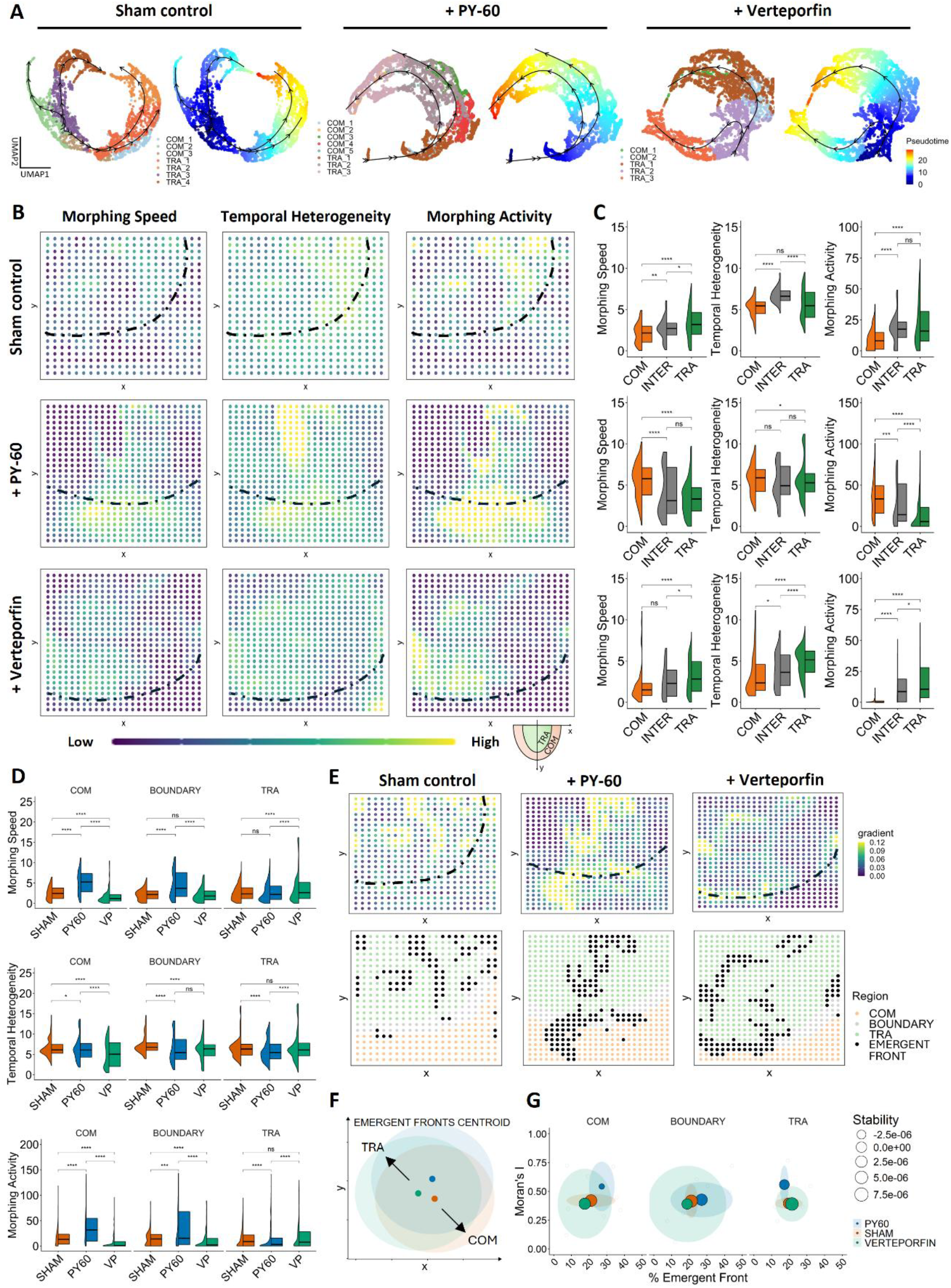
Kinemomics resolved spatiotemporal morphing dynamics in distinct myocardial layers during ventricular morphogenesis. **A)** High-dimensional omics approaches based on spatiotemporal resolved local cellular kinematics changes over 3 hr live tracking. Trajectory inference based on actual cell tracking time-derived pseudotime, which represents the morphing state of collective cell kinematics across developmental stages. **B)** Spatial mapping of local cellular morphing speed, d(pseudotime)/d(tracking time); temporal heterogeneity, sd(pseudotime); and morphing activity, morphing velocity * heterogeneity. **C-D)** YAP activation led to stronger morphing activity in the compact and intermediate boundary zone between compact and trabecular myocardium, whereas YAP inhibition resulted in higher morphing activity in the trabecular region. **E)** Spatial map of morphing activity gradient (top). Position with high gradient magnitude (0.8 quantile) indicates sharp morphing activity changes and was defined as emergent fronts (bottom). **F)** YAP activity alteration led to shifting of the centroids of emergent fronts. **G)** Moran’s Index – emergent fronts proportion, with circle size indicating front stability. *p <= 0.05, **p <= 0.01, ***p <= 0.001,****p <= 0.0001 determined via pairwise Mann–Whitney U test with biological replicates N=3 for each condition.

YAP perturbation markedly altered these patterns. YAP activation increased morphing activity in the compact layer, driven by elevated morphing speed and temporal heterogeneity, while suppressing morphing activity in the trabeculae. Conversely, YAP inhibition reduced morphing speed and temporal heterogeneity within the compact layer, resulting in markedly weakened morphing activity (Fig. 6C-D). Overall, YAP GOF enhanced morphing activity in the compact and intermediate regions by accelerating local morphogenetic progression, whereas YAP LOF reduced morphing activity through reduced progression and heterogeneity (Fig. 6D). These results were consistent with earlier observations that YAP inhibition prolonged compact cardiomyocyte calcium transient duration, shortened ECC response, and suppressed proliferation (Fig. 4A-C), reflecting a shift toward a more functionally mature but less dynamically remodeling state^52^. Collectively, these findings highlight the power of spatiotemporal Kinemomics not only to identify distinct treatment- and time-specific phenotypes, but also to resolve their spatiotemporal evolution.

### Kinemomics time-resolved phenotype evolution uncovers emergent developmental fronts

Finally, we examined whether spatiotemporal Kinemomics could identify localized morphogenetic fronts driving collective tissue morphology. Analogous to our Kine-Bio Multi-Omics framework, we computed spatial gradients of morphing activity and defined the top 20% of gradient magnitudes as emergent developmental fronts within each sample (Fig. 6E, S11). These fronts were detected across the myocardial wall under all conditions, but their spatial centroids shifted toward trabecular tips and the ventricular lumen following YAP perturbations (Fig. 6F). We further quantified front dynamics using the divergence of morphing activity as a measure of stability and Moran’s I as an index of spatial autocorrelation^53,54^. YAP activation increased spatial coherence of emergent fronts in both compact and trabecular layers, as reflected by higher Moran’s I, while simultaneously increasing divergence and reducing stability in the compact layer (Fig. 6G). In contrast, YAP LOF enhanced front convergence and stability within trabeculae while modestly reducing emergent fronts in the compact and intermediate boundary regions. Together, these findings suggest that the compact myocardium is more sensitive to YAP-dependent morphogenetic regulation, whereas alternative signaling programs may predominate in trabecular remodeling under YAP LOF.

## Discussion

Organotypic brain slice culture has been widely used in neurodevelopmental studies^55,56^. Similar approaches in adult myocardium have also enabled long-term culture while preserving cardiomyocyte contractility under external mechanical stimulation for electrophysiological, pathophysiological, and pharmacological studies in both healthy and diseased tissue^57,58^. Here, we introduce a novel fetal cardiac organotypic slice culture system that preserves both beating and morphogenesis, enabling controlled perturbations and real-time analysis of mesoscale myocardial morphodynamics, including cell migration, tissue morphology, and contractile dynamics. These live slices maintain autonomous calcium handling without external stimulation, enabling assessment of ECC within a physiologically heterogeneous full-thickness myocardium. In addition, direct monitoring of cellular migration and tissue morphology reveals dynamic cell-tissue kinematic interactions underlying fetal growth and remodeling, extending beyond static snapshot-based analyses.

Omics approaches, including genomic, transcriptomic, proteomic, and metabolomic profiling, have enabled high-dimensional phenotyping of cell populations. Concurrently, multi-omics pipelines integrating transcriptional data with mechanical stress^59^, immunofluorescence imaging^60^, and metabolomic and lipidomic profiling^61^, have begun to bridge molecular and tissue-scale processes, highlighting cross-scale interactions. However, these largely static atlases remained limited in resolving morphodynamic behaviors during development^8,9,62^. Time-resolved analyses of cellular morphology and metabolism from live imaging often rely on precise single-cell segmentation^63–66^, which remains challenging in dense three-dimensional tissues with irregular cell shapes, such as the myocardium. Building on these insights, we instead derive spatiotemporally resolved mesoscale kinematics using a finite-difference framework, eliminating the need for single-cell segmentation that requires extensive model training and precise membrane labeling. Spatial resolution can be flexibly tuned by adjusting grid size, enabling adaptation to imaging trade-offs between penetration depth, acquisition speed, and resolution in 3D systems. Importantly, Kinemomics extends morphodynamic analysis to intact, full-thickness myocardial tissue.

By incorporating endpoint assays targeting mechanosensitive pathways, we further establish a spatiotemporal multi-omics pipeline that co-registers cellular kinematics with biomolecular activity to reconstruct high-dimensional morphodynamic profiles. This framework resolves layer-, treatment-, and time-dependent phenotypes, identifies emergent or maladaptive developmental trajectories and fronts driving system behavior, and reduces computational complexity while bypassing reliance on transcriptional data. It enables faster, more cost-effective, and higher-dimensional characterization of disease progression and critical transition thresholds. Beyond cardiac development, this framework is readily extensible to other morphodynamic processes, including cancer invasion and metastasis, immune cell trafficking, and tissue regeneration, with potential to inform diagnostic and therapeutic strategies targeting cellular decision-making at critical spatiotemporal junctures. More broadly, it provides a generalizable foundation for computational morphodynamics in complex systems governed by cross-scale interactions.

## Methods

### Chick embryo slice culture and preparation

Fertilized white Leghorn eggs were incubated blunt end up in a continuous rocking incubator at 37.5°C and approximately 50% humidity until the desired stage. On the 7th day after incubation, embryos were euthanized by decapitation in ice-cold HBSS (Gibco, 14025092) under a dissection microscope. Hearts were isolated from the embryos and rapidly embedded in an agarose block. The agarose block was made with 4% low-melting-point agarose (VWR, 97062-398) in Medium 199 (Sigma-Aldrich, M2154) and kept at approximately 40°C during embedding. Hearts were embedded in frontal orientation and sectioned into 200-250 μm thickness slices along the long axis using a vibrating microtome. After sectioning, selected heart slices with four-chambered view were cultured in Medium 199 with 10% chicken serum (Gibco, 16110082) and 1% Penicillin-Streptomycin (Gibco, 15140122) with ambient oxygen and 5% CO2 at 37°C for at least 1–2 hr for recovery. Only slices without arrythmia were used for imaging. Physiological circumferential stress was maintained through compressive strain loaded via a permeable membrane. The strain can be modulated by adjusting the height between the permeable Polydimethylsiloxane (PMDS) membrane to the bottom of culture dish, which controls the compressed thickness of heart slices (Supplementary Note 1). Strain loaded on the slice was designed based on mean wall stress at *in vivo* diastolic pressure at Hamburger-Hamilton (HH) stage 30 ^67,68^. All experimental procedures were approved by Cornell University Institutional Animal Care and Use Committee (IACUC).

### Calcium imaging and analysis

Calbryte 520 AM (AAT Bioquest, 20651) was used as the fluorescent calcium indicator.

Slices were incubated with 10 μM Calbryte 520 AM in Medium 199 with 0.04% Pluronic F-127 (Sigma-Aldrich, P2443-250G) for 1–2 hr in CO2 incubator prior to imaging. In alternative treatments, 5mM caffeine (Abcam, AB120240), 10μM epinephrine (Abcam, AB142299), 3µM verapamil (Thermo Scientific, 32933-0010), 10μM PY-60 (MedChemExpress, HY-141644), or 10μM verteporfin (Sigma-Aldrich, SML0534) were used together with Calbryte 520 AM during incubation. After dye loading, slices were transferred into glass-bottom confocal dish and imaged in Medium 199 with 25mM HEPES (Sigma-Aldrich, M7528) and 10% chick serum. Imaging was performed with a Zeiss LSM710, where calcium transient images were acquired with a x10, 0.45 numerical aperture (NA) water objective (Zeiss) with x4 zoomed in and 128×128 pixels and a frame per second of 20-30.

Cardiac contraction in slice culture was approximated into locally linear movement within the 2D plane, thus rigid-body registration was performed to remove any contraction induced calcium signal changes using StackReg^69^. Calcium traces were measured based on signal intensity using FIJI ImageJ. Calcium analysis was performed using MATLAB (The MathWorks Inc., Natick, USA) based program. Measured calcium traces were normalized to baseline, which was defined as the mean value of the lower 10% of calcium, and averaged into one full calcium trace. The following parameters were defined and analyzed: 1) maximum normalized amplitudes (Fmax/F0) as maximum fluorescence value (Fmax) divided by baseline value (F0); 2) beating frequency as the reciprocal of the average time interval between two adjacent peaks; 3) transient duration (CaD50) as the time duration during which calcium is above 50% of the signal magnitude; 4) rise time (T50 on) as the time from 50% activation to peak; 5) time of off-reaction (T50 off) as the time from peak to 50% relaxation; and 6) the time constant of decay (Tau).

For excitation-contraction coupling, cellular contraction trace was measured as displacement from rest position to contraction position at each time point within a full contraction cycle. Calcium transient tracing *F*(*t*)/*F*_0_ and contraction tracing *du*(*t*) were fitted into Gaussian equations for data interpolation as below:

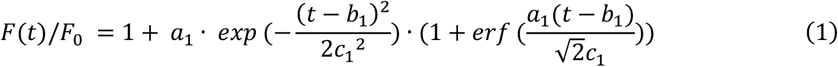

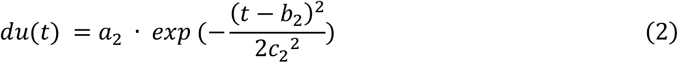

The time difference between two peaks was then taken as the ECC time.

### Timelapse imaging and cell tracking

For timelapse imaging, Calcein AM (Invitrogen, C3100MP) was used for cell labeling and cell vitality monitoring. Slices were incubated with 10 μM Calcein AM for 1–2 hr in CO2 incubator prior to imaging. For YAP perturbation study, slices were treated with 10 μM PY-60 (MedChemExpress, HY-141644) or 10 μM verteporfin (Sigma-Aldrich, SML0534) together with Calcein AM. Heart slices were then transferred into glass-bottom confocal dish and imaged in Medium 199 with 25 mM HEPES and 10% chick serum. Cell migrations were followed longitudinally for up to 4 hours via fluorescence microscopy with a Zeiss LSM710. Z-stack images were taken every 10 minutes with each scan time controlled under 3 minutes to minimize photobleaching. After imaging, slices were fixed in 4% Paraformaldehyde (PFA; Sigma-Aldrich, 252549) at 4°C overnight to prepare for further endpoint assay.

The timelapse images were projected in z-axis with maximum intensity, and all cell movements were assumed to be within the 2D plane. Translational and rotation registration was performed using StackReg to eliminate global tissue sliding or shifting during prolonged timelapse imaging. Cell tracking and local tissue tracking were then performed using TrackMate Kalman tracker and Particle Image Velocimetry (PIV) as FIJI ImageJ plugins^70–73^, respectively. For each cell tracked, spatiotemporal features including spatial position of the cell (x, y, t), displacements in x and y directions (dx, dy, t), speed, track linearity, and directional change rates were detected and saved for further analysis.

### Endpoint immunofluorescence staining

After live imaging, heart slice sections were fixed with 4% PFA overnight at 4°C, washed with 1x phosphate-buffered saline (PBS; Gibco, 20012-027) twice for at least 1 hr, placed in 15% sucrose (Thermo Scientific, A15583.36) in PBS at 4°C until sinks, then transferred into 30% sucrose in PBS until sinks. Heart slices were then placed into cryomolds (Tissue-Tek) and embedded in O.C.T. Compound (Tissue-Tek) with chambered view facing to the base. Tissue was frozen in the O.C.T-filled mold using liquid nitrogen and stored at −80°C until further usage. For immunofluorescence staining, daughter sections of 10-20 μm were generated using a cryostat. Tissue sections were permeabilized with 0.3% Triton X-100 (VWR, 01279-1L) and 0.3 M glycine (VWR, 0167-1KG) in PBS for 30 min and incubated in a blocking solution containing 0.3% Triton X-100, 0.3 M glycine, 5% goat serum (CELLect, 2939149), and 1% bovine serum albumin (BSA; Sigma-Aldrich, A9647-100G) in PBS for 1 hr. Samples were then incubated with primary antibodies diluted in PBS with 0.2% Tween-20 (VWR, 0777-1L) and 1% BSA (1% BSA in 0.2% PBST) overnight at 4°C, followed by five washes in 1% BSA in 0.2% PBST. Secondary antibodies and nucleus counterstain were then incubated in 1% BSA in 0.2% PBST for 1 hr at room temperature, followed by five washes in PBS. Slides were then mounted with Prolong Glass Antifade Mountant (Invitrogen, P36984) and stored at room temperature in dark until microscope imaging. The source and dilution ratio of primary and secondary antibodies used in this study are listed in Supplementary Table 1.

### Feature extraction and normalization

Staining images were taken at approximately the same location as the live cell tracked region. Images were then meshed into 16×16 pixel (0.2768 micron/pixel) square grid of sampling element to cover the entire field of image. This resulted in 6,000 – 10,000 elements per biological replicate. Normalized mean fluorescent intensities of each channel were calculated for each element. To achieve local continuity, intensities were averaged among the neighboring 16 elements and stored as the local mean intensity of the central element. A custom macro in FIJI ImageJ was used to extract the x and y position of element centroid and the corresponding local mean intensities for further analysis. Same grid elements were spatially mapped to PIV cell tracking results. Local mean displacements, migration directions, and fluorescent intensities of biomolecules were then stored in the same x and y position for spatial analysis. Kine-Bio co-mapping was only performed for cell tracking results from the start of imaging to the end of imaging without intermediate temporal information included. For spatiotemporal Kinemomics, local cellular kinematics was resolved with 30 minutes temporal intervals, with grid spacing increased to 64 pixels to reduce computational overhead. Local fluorescence intensities extracted from the Calcein AM and nucleus counterstain channels for each sample in live timelapse imaging and endpoint immunofluorescence staining, respectively, were used to determine the void space between trabeculae. Sampling grids with a mean Calcein AM or nucleus intensity smaller than 0.1 were excluded for omics analysis.

### UMAP and clustering

UMAP dimensionality reduction was performed using the R package umap (v0.2.10.0)^74^. Gaussian mixture model (GMM) clustering was performed with a varying number of clusters (from 2 to 10) using the R package gmm (v1.9-1)^75^. Bayesian Information Criterion (BIC) and silhouette scores were computed for each cluster to determine an optimal user-defined number of clusters. For spatial Kinemomics, only kinematic features from start of imaging to end of imaging were used for UMAP and clustering. All samples of the same treatments were pooled together. Technical variation correction was performed using Harmony (v1.2.4)^76,77^ in R for each feature extracted across multiple samples of the same treatment to minimize batch effects. Results were visualized spatially, where each point is colored according to its associated cluster. A consensus spatial map was generated for each treatment, showing the most likely cluster at each point across all samples based on frequency. For Kine-Bio Multi-Omics across-treatments, GMM was performed within distinct myocardial layer groups based on ground truth. Spatial mapping was then performed for each sample according to the cell’s native coordinates. For spatiotemporal Kinemomics, UMAP and clustering were performed separately for each sample to reveal individual morphological trajectories.

### Temporal evolution and pseudotime analysis

For trajectory inference and pseudotime analysis, R package slingshot (v 2.14.0) was used^7^. In Kine-Bio Multi-Omics across treatments, a binary feature was assigned to each point in sham condition with 0, whereas points in treated conditions were assigned as 1, representing malforming score. The cluster with the minimum mean malforming score was automatically selected as the start cluster within each region. Pseudotime was then calculated based on malforming score and mapped spatially, representing malforming progression. Local least-square gradient method was performed to estimate the spatial gradient of malforming progression as below:

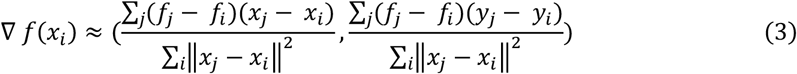

where *f*_*i*_ is gradient of malforming progression at point *i, f*_*j*_ is the neighbor values. Points with a malforming gradient magnitude larger than 0.45 were identified as the malforming fronts. In spatiotemporal Kinemomics, pseudotime analysis was performed based on recorded imaging time (tracking time), representing morphing progression. The cluster with the shortest mean tracking duration was automatically selected as the start cluster. The slope of pseudotime – imaging time was taken as the morphing speed, and the standard deviation of the pseudotime was taken as morphing temporal heterogeneity. Product of morphing speed and temporal heterogeneity was identified as morphing activity, which represents how fast and dynamic the morphology is at each spatial location. Local least-square gradient method was used to estimate the spatial gradient of activity using the same equation (3). Points exceeding the 0.8 quantile of activity gradient magnitudes were identified as the emergent developing fronts.

Further, we calculated distance-weighted discrete Laplacian of the scalar field morphing activity to determine the divergence of gradient for each point *i*:

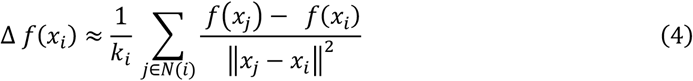

where *f*(*x*_*i*_) is activity at point *i, N*(*i*) is neighbors within a selected radius, *k*_*i*_ is the number of neighbors. Stability of the identified emergent fronts was defined as the negative mean product of gradient magnitude and divergence as below:

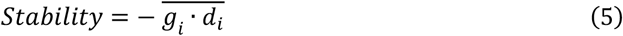

where *g*_*i*_ and *d*_*i*_ is the activity gradient magnitude and divergence at point *i*, respectively. A large positive stability value then indicates stable fronts with strong convergence, while a large negative value suggests unstable fronts exhibiting a strong divergence.

In addition, Moran’s I was computed using the following formula to analyze the spatial autocorrelation of fronts:

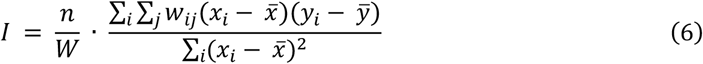

where *n* is the number of observations, *w*_*ij*_ = 1/*k*_*i*_ is the spatial weights, *k*_*i*_ is the number of neighbors of point *i*, then *W* = ∑_*i*_ ∑_*j*_ *w*_*ij*_ = *n* is the sum of all weights.

## Statistical analysis

Statistical comparisons were made using R package ggplot2 (v3.4.4)^78^. All statistics were determined via t-tests, ANOVA, or pairwise Mann–Whitney U test. P<0.05 denotes significance.

## Data availability

All data and analysis are available to be shared upon request.

## Code availability

Packages and software used for this study are open source. Custom MATLAB codes and FIJI ImageJ macros are available upon request.

## Acknowledgement

This work was supported by National Institutes of Health (R01 HL160028 to J.T.B.), National Science Foundation (EF-2222434 to J.T.B.), and Additional Ventures Society (to J.T.B.). Imaging data was acquired through the Cornell Institute of Biotechnology’s Imaging Facility, with NIH 1S10RR025502 funding for the shared Zeiss LSM 710 Confocal Microscope.

The authors acknowledge the support of Warren Zipfel and Alex Kwan for generously sharing vibratome, and Adrienne Roeder for Confocal Microscope. The authors also thank Mong Lung Steve Poon and Lingxin Meng for invaluable suggestions on figure panel design and spatial analysis methods, respectively.

## Author contribution

Conceptualization: G.D. and J.T.B.; Methodology: G.D., L.W., D.M., M.W., J.T.B.; Experimental investigation: G.D., L.W., J.C.; Data analysis: G.D., L.W., J.C.; Software: G.D. and L.W.; Data interpretation: G.D. and L.W.; Visualization: G.D.; Resources: J.T.B.; Writing - original draft: G.D. and L.W.; Writing - review & editing: J.T.B.; Supervision: J.T.B.; Project administration: J.T.B.; Funding acquisition: J.T.B.

